# A potent neutralizing human antibody reveals the N-terminal domain of the Spike protein of SARS-CoV-2 as a site of vulnerability

**DOI:** 10.1101/2020.05.08.083964

**Authors:** Xiangyang Chi, Renhong Yan, Jun Zhang, Guanying Zhang, Yuanyuan Zhang, Meng Hao, Zhe Zhang, Pengfei Fan, Yunzhu Dong, Yilong Yang, Zhengshan Chen, Yingying Guo, Jinlong Zhang, Yaning Li, Xiaohong Song, Yi Chen, Lu Xia, Ling Fu, Lihua Hou, Junjie Xu, Changming Yu, Jianmin Li, Qiang Zhou, Wei Chen

**Affiliations:** Beijing Institute of Biotechnology, Academy of Military Medical Sciences (AMMS), Beijing 100071, China; Key Laboratory of Structural Biology of Zhejiang Province, Institute of Biology, Westlake Institute for Advanced Study, School of Life Sciences, Westlake University, Hangzhou 310024, Zhejiang Province, China; Beijing Advanced Innovation Center for Structural Biology, Tsinghua-Peking Joint Center for Life Sciences, School of Life Sciences, Tsinghua University, Beijing 100084, China

## Abstract

The pandemic of coronavirus disease 2019 (COVID-19) caused by severe acute respiratory syndrome coronavirus 2 (SARS-CoV-2) presents a global public health threat. Most research on therapeutics against SARS-CoV-2 focused on the receptor binding domain (RBD) of the Spike (S) protein, whereas the vulnerable epitopes and functional mechanism of non-RBD regions are poorly understood. Here we isolated and characterized monoclonal antibodies (mAbs) derived from convalescent COVID-19 patients. An mAb targeting the N-terminal domain (NTD) of the SARS-CoV-2 S protein, named 4A8, exhibits high neutralization potency against both authentic and pseudotyped SARS-CoV-2, although it does not block the interaction between angiotensin-converting enzyme 2 (ACE2) receptor and S protein. The cryo-EM structure of the SARS-CoV-2 S protein in complex with 4A8 has been determined to an overall resolution of 3.1 Angstrom and local resolution of 3.4 Angstrom for the 4A8-NTD interface, revealing detailed interactions between the NTD and 4A8. Our functional and structural characterizations discover a new vulnerable epitope of the S protein and identify promising neutralizing mAbs as potential clinical therapy for COVID-19.

---

The global outbreak of COVID-19 has emerged as a severe threat to human health (*1–4*), claiming over two hundred and sixty thousand lives as 7 May, 2020 (*5*). COVID-19 is caused by a novel coronavirus, the severe acute respiratory syndrome coronavirus 2 (SARS-CoV-2), which is an enveloped, positive-strand RNA virus that causes upper respiratory diseases, fever and severe pneumonia in humans (*2, 3, 6*).

SARS-CoV-2 is a new member of the β coronavirus genus, which also contains SARS-CoV and MERS-CoV that caused epidemic in 2002 and 2012, respectively (*7, 8*). SARS-CoV-2 shares about 80% sequence identity to SARS-CoV, implying similar infection mechanism (*2*). Indeed, SARS-CoV-2, same as SARS-CoV, hijacks angiotensin-converting enzyme 2 (ACE2) as cellular receptor(*9–19*).

The trimeric S protein covers the surface of coronavirus and plays a pivotal role during viral entry (*20, 21*). The S protein is cleaved into the N-terminal S1 subunit and C-terminal S2 subunit by host proteases such as TMPRSS2 (*21, 22*) and undergoes conformational change from prefusion to postfusion state during infection (*23*). S1 and S2 mediate receptor binding and membrane fusion, respectively (*18*). S1, consisting of the N-terminal domain (NTD) and the receptor binding domain (RBD), is critical in determining tissue tropism and host ranges (*24, 25*). The RBD is responsible for binding to ACE2, while the function of NTD is not well understood. In some coronaviruses, the NTD may recognize specific sugar moieties upon initial attachment and might play an important role in the prefusion to postfusion transition of the S protein (*26–29*). The NTD of the MERS-CoV S protein can serve as a critical epitope for antibody neutralizing (*29*).

The SARS-CoV-2 S protein-targeting monoclonal antibodies (mAbs) with potent neutralizing activity have become the focus of therapeutic interventions for COVID-19 (*30–32*). Most SARS-CoV-2 antibodies reported previously were sorted to target the RBD in order to inhibit the association between the S protein and ACE2 (*31–34*). The RBD-targeting antibodies, when applied individually, may induce viral escaping mutations, thus fostering evolution of new viral strains that are insensitive to these antibodies (*29*). Antibodies targeting non-RBD regions may serve as “cocktail” therapeutics for SARS-CoV-2.

Here we report the isolation and characterization of S protein-specific monoclonal antibodies derived from memory B or plasma B cells of COVID-19 survivors. We also determined the structure of the complex between the SARS-CoV-2 S protein and 4A8, one of the isolated mAbs, at an overall resolution of 3.1 Å and local resolution of 3.4 Å for the interfaces between 4A8 and the S protein. 4A8 targets the S-NTD with potent neutralizing activity. These findings reveal a new epitope of the SARS-CoV-2 S protein for antibody neutralizing, which may lay the foundation for new prophylactic and therapeutic interventions for SARS-CoV-2.

## Results

### Isolation of human mAbs from memory B cells and plasma B cells

To isolate monoclonal antibodies and analyze the humoral antibody responses to SARS-CoV-2, we collected plasma and peripheral blood mononuclear cells (PBMCs) from 10 patients recovered from SARS-CoV-2 infection. The age of donors ranges from 25 to 53 years. The interval from blood collection date to disease confirmation date ranged from 23 to 29 days for No. 1-5 patients and 10 to 15 days for No. 6-10 patients (Table S1). We evaluated the titers of binding antibodies in plasma to different fragments of the SARS-CoV-2 S protein, including the extracellular domain (S-ECD), S1, S2, and RBD, and the Nucleocapsid (N) Protein. Plasma from all the patients except donor No. 2 bound to all 5 SARS-CoV-2 protein segments, while that from donor No. 2 recognized S-ECD and S2 only (Fig. 1A). The neutralizing capacities of plasma against live SARS-CoV-2 and HIV-vectored pseudotyped SARS-CoV-2 are correlated (r=0.6868, p<0.05) (Fig. 1B). These results indicate that humoral immune responses were specifically elicited for all of the 10 patients during the natural infection of SARS-CoV-2.

**Fig. 1.**
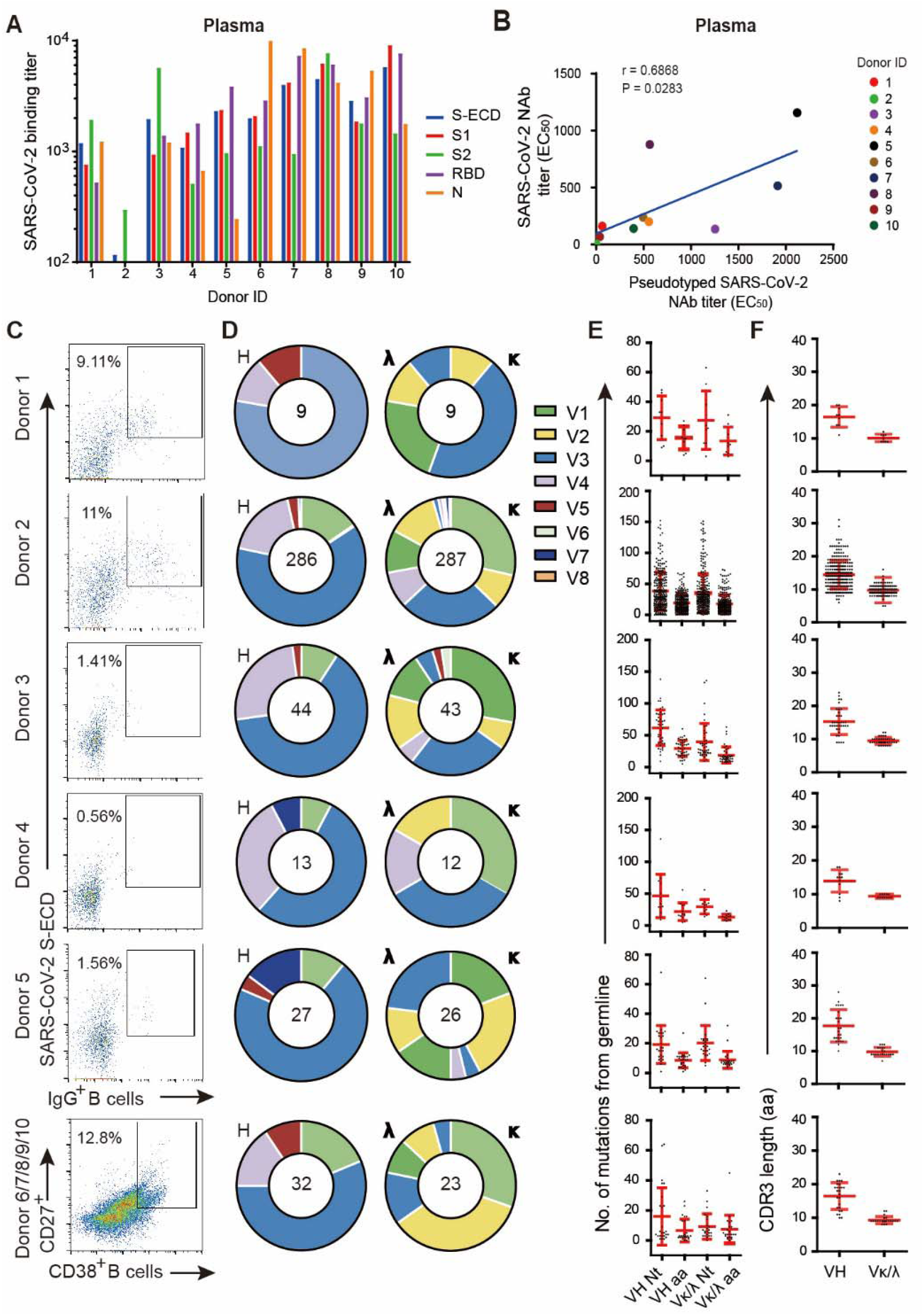
Isolation of antigen-specific monoclonal antibodies from convalescent patients of SARS-CoV-2. **(A**) Reactions of plasma to SARS-CoV-2 proteins. S-ECD (extracellular domain of S protein), S1, S2, RBD (receptor binding domain) and N (nucleotide protein) were used in ELISA to test the binding of plasma. Plasma of heathy donors were used as control, and cut-off values were calculated as O.D. 450 of control x 2.1. (**B**) The correlations between the authentic SARS-CoV-2 neutralizing antibody titers and the pseudotyped SARS-CoV-2 neutralizing antibody titers in plasma. Neutralizing assays of plasma against authentic SARS-CoV-2 were performed using Vero E6 cells, and neutralization against pseudotyped SARS-CoV-2 were determined using ACE2-293T cells. The correlations were calculated by Pearson correlation test in Graphpad 7.0. **(C)** Flow cytometry sorting from PBMCs of 10 convalescent patients. **(D)** Distribution of V gene families in heavy and light chains of all unique clones (the total number is shown in the center of the pie charts) for each donor. **(E)** The number of mutations from the germline of all clonal sequences identified in (**D**) was shown. **(F)** CDR3 amino acid lengths of VH and VL of all clonal sequences identified in (**D**).

To isolate S protein-specific monoclonal antibodies, we first sorted the IgG^+^ memory B cells from peripheral blood mononuclear cells (PBMCs) of the No. 1-5 convalescent patients with flow cytometry using S-ECD as probe (Fig. 1C). The percentage of S-ECD-reactive IgG^+^ B cells ranges from 0.56% to 11% as revealed by fluorescence activating cell sorter (FACS). To avoid bias introduced by S-ECD, we sorted plasma B cells from mixed PBMCs derived from another five convalescent patients (No. 6-10) without any antigen-specific probes. The percentage of plasma B cells in CD3-CD19^+^ B cells was 12.8%, higher than that of memory B cells (Fig. 1C).

From the sorted B cells, we identified 9, 286, 43, 12 and 26 clones of single B cell from patients No. 1 to 5, respectively, and 23 clones of single B cell from the mixed PBMCs of patients No. 6 to 10 (Fig. 1D). The distribution of the sequenced heavy (IgH) gene families was comparable among the 10 donors, with VH3 being the most commonly used VH gene, while different donors displayed variable preferences for the light chain (IgL) gene families (Fig. 1D). The combination of V3 and J4, V3 and D3, and D3 and J4 were the most common usage for the IgH gene family (Fig. S1). The plasma B cells have less mutations than memory B cells in both heavy variable chains (VH) and light variable chains (VL) (Fig. 1E), suggesting lower levels of somatic hypermutation (SHM) in plasma B cells than memory B cells after SARS-CoV-2 infection. The lengths of complementarity-determining region (CDR) 3 for antibodies were similar among the donors, ranging from 13.9 to 17.7 for VH and 9.3 to 10.1 for VL (Fig. 1F).

### Binding profiles of SARS-CoV-2 S protein-specific human mAbs

To screen S protein-specific antibodies, we determined the binding specificity using ELISA for the 399 human mAbs sorted above. 1, 16, 1, 3, and 9 S-ECD-specific mAbs were identified from donors No. 1-5, respectively. A total of 5 mAbs were identified from donors No. 6-10 (Fig. 2A). The sequence identities of CDRH3 in the 35 S-ECD-specific mAb ranged from 40.9% to 97.6% (Fig. S2 and Table S2). We further characterized domain specificities of the 35 mAbs with different fragments of the S protein, including S1, S2 and RBD (Fig. 2A). The S-reactive mAbs are classified into 4 major groups based on their EC_50_ values (Fig. 2A). Group 1 recognizes only S-ECD. Group 2 recognizes S-ECD and S1, with subgroup 2A binding S-ECD and S1 and subgroup 2B binding S-ECD, S1, and RBD. Group 3 interacts with both S1 and S2, with subgroup 3A targeting RBD and subgroup 3B fails to bind RBD. Group 4 recognizes S-ECD and S2. Remarkably, only 4 mAbs recognize RBD among the 35 S-specific mAbs (Figure 2A and 2B).

**Fig. 2.**
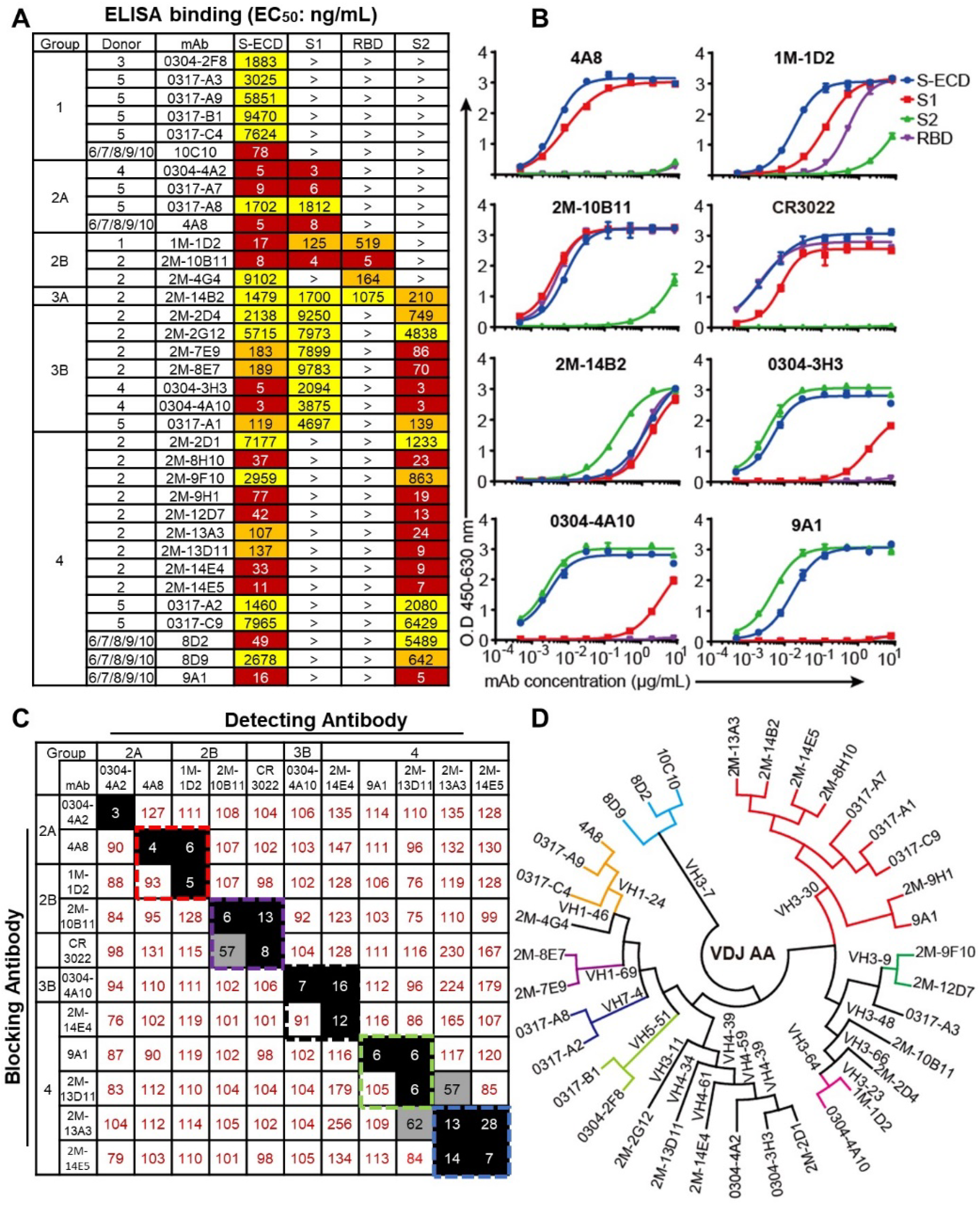
Binding profiles of Spike protein-specific mAbs. **(A)** Heatmap showing the binding of mAbs to different types of spike proteins determined using ELISA. The EC_50_ value for each S-mAb combination is shown, with dark red, orange, yellow, or white shading indicating high, intermediate, low, or no detectable binding, respectively. EC_50_ values greater than 10,000 ng/ml are indicated (>). **(B)** Binding curves of representative mAbs. CR3022 is a control that was reported to bind SARS-CoV and SARS-CoV-2 RBD. **(C)** Heatmap showing the competing binding of some representative S-reactive mAbs assayed in ELISA. Numbers in the box indicate the percentage binding of detecting mAb in the presence of the blocking antibody compared to the binding of detecting mAb in the absence of the blocking antibody. The mAbs were considered competing if the inhibiting percentage is <30% (black boxes with white numbers). The mAbs were judged to non-compete for the same site if the percentage is >70% (white boxes with red numbers. Gray boxes with black numbers indicate an intermediate phenotype (30%~70%). **(D)** Phylogenetic trees of all the S-specific mAbs.

A competition-binding assay using ELISA were performed for several representative mAbs to determine if there are any overlapping antigenic sites within distinct binding groups or between different mAbs, with CR3022 being used as a control that cross-reacts with the RBD of both SARS-CoV and SARS-CoV-2 (*35*) (Fig. 2C). Among these mAbs, 4A8 competed with 1D2. Another RBD-reactive mAb, 2M-10B11, competed with CR3022, suggesting overlapped epitopes on RBD for these two mAbs. These results indicate that the S-specific antibodies elicited by SARS-CoV-2 infection target at least four antigenic regions on the S protein of SARS-CoV-2.

To characterize the diversity in gene usage and affinity maturation, the phylogenetic trees of these S-specific mAbs were analyzed based on the amino acid sequences of VHDJH and VLJL using a neighbor-joining method in MEGA7 Software. Results indicate that the VH gene usage is very diverse among the 35 mAbs from 10 donors, with VH 3-30 being the most frequently used germline gene. Moreover, there was no particularly favored VH gene identified among S1, S2, or RBD-reactive mAbs (Fig. 2D).

### Neutralizing activities of SARS-CoV-2 S-specific human mAbs

We first performed *in vitro* neutralization studies of the 35 S-specific mAbs using live SARS-CoV-2 virus in Vero-E6 cells (Fig. 3A). 1M-1D2 and 4A8 exhibited medium to high neutralizing capacity with EC_50_ of 23.99 and 0.607 μg/ml, respectively, while the neutralizing curves of 0304-3H3 cannot be fit well. As expected, the RBD-targeting control mAb, CR3022, failed to neutralize live SARS-CoV-2 (*35*). Moreover, while the CR3022-competing mAb, 10B11, bound to the SARS-CoV-2 RBD with EC_50_ of 5 ng/ml, it also failed to neutralize live SARS-CoV-2. These results suggest that binding affinities of mAbs against RBD did not correlate fully with the neutralizing abilities of mAbs. To further investigate the inhibitory activity of 4A8, 0304-3H3, and 1M-1D2 to live virus, we tested the RNA load of live viruses in Vero-E6 cells treated with each mAb using real time qPCR (Fig. 3B). Consistent with the cytopathic effect (CPE) assay results (Fig. 3A), 4A8 displayed higher inhibitory capacities than 1M-1D2.

**Fig. 3.**
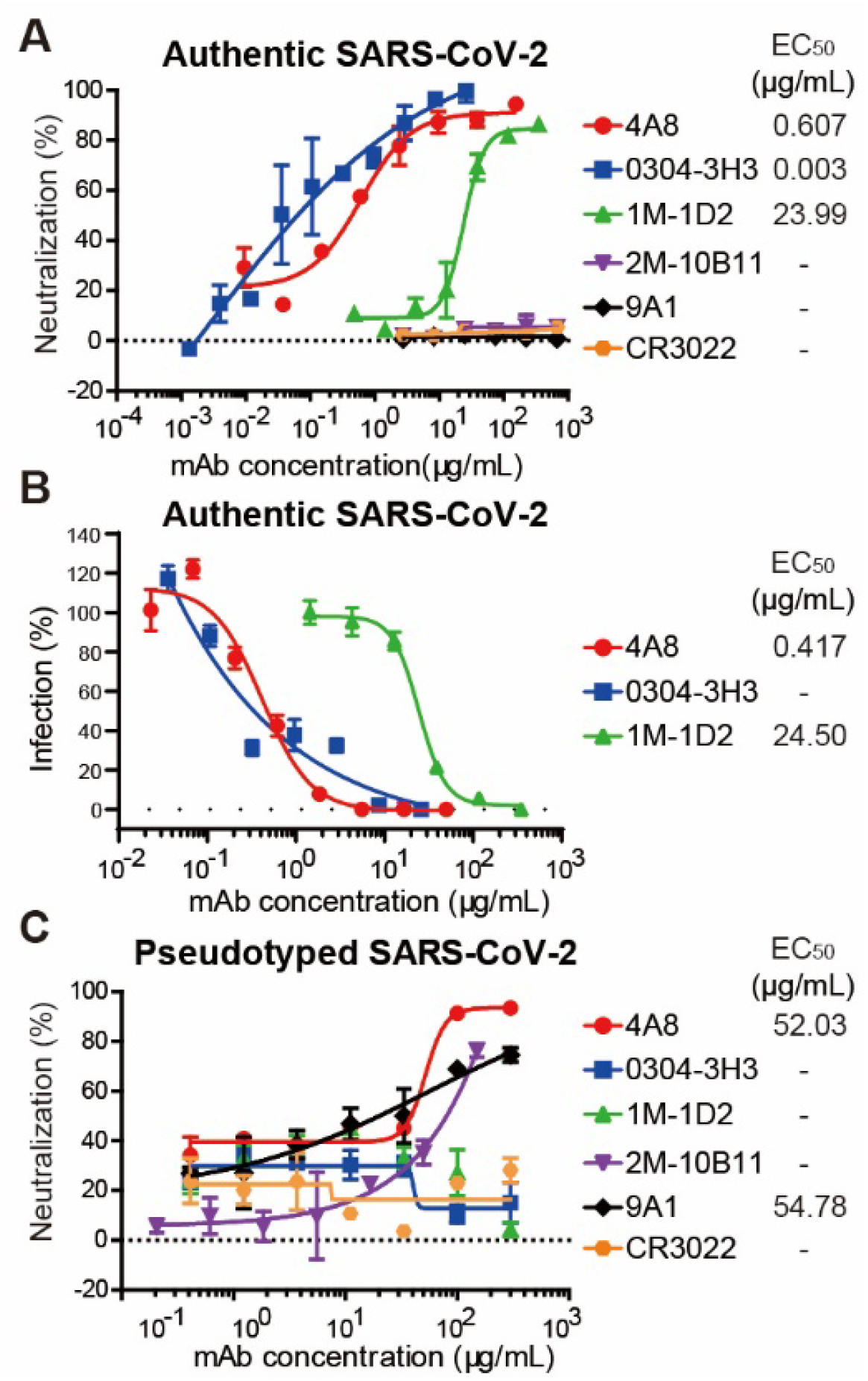
Neutralizing capacities of S-reactive mAbs. **(A)** Neutralization of S-reactive mAbs to authentic SARS-CoV-2 in Vero-E6 cells. **(B)** The authentic SARS-CoV-2 virus RNA load was determined in Vero-E6 cells treated with S-reactive mAbs using qPCR. Percent infection was calculated as the ratio of RNA load in mAb-treated wells to that in wells containing virus only. **(C)** Neutralization of S-reactive mAbs against HIV-vectored pseudotyped SARS-CoV-2 in ACE2-293T cells. data were shown as mean ± SD of a representative experiments.

We next performed luciferase reporter gene assays for all 35 S-binding mAbs using HIV-vectored pseudotyped SARS-CoV-2, among which 3 mAbs exhibited neutralizing activity against the pseudotyped virus (Fig. 3C). 4A8 protected ACE2-293T cells with EC_50_ of 52.03 μg/ml, while neutralization of 0304-3H3 and 1M-1D2 were not observed. 10B11 and 9A1 showed a minor level of inhibition. These findings suggest that the results of pseudotyped SARS-CoV-2 were not entirely consistent with that of live SARS-CoV-2. This difference is likely explained by the insusceptibility of pseudotyped SARS-CoV-2 to some mAbs with specific neutralizing mechanisms. 4A8 is likely to be a potential candidate for the treatment of SARS-CoV-2, since 4A8 displayed high levels neutralizing capacities against both authentic and pseudotyped SARS-CoV-2.

### Binding characterization of candidate mAbs

To determine the possible neutralizing mechanism of the mAbs, we first determined the binding affinities of mAbs to different segments of the S protein, including S-ECD, S1, S2, and RBD using bio-layer interferometry. All 5 tested mAbs bound to S-ECD with high affinity, with the equilibrium dissociation constant (K_D_) less than 2.14 nM (Fig. 4A). 4A8, 1M-1D2, and 10B11 bound to S1 with similar K_D_ ranging from 92.7 to 159 nM, whereas 0304-3H3 and 9A1 targeted S2. Additionally, 10B11 were shown to bind RBD with K_D_ of 26.8 nM.

**Fig. 4.**
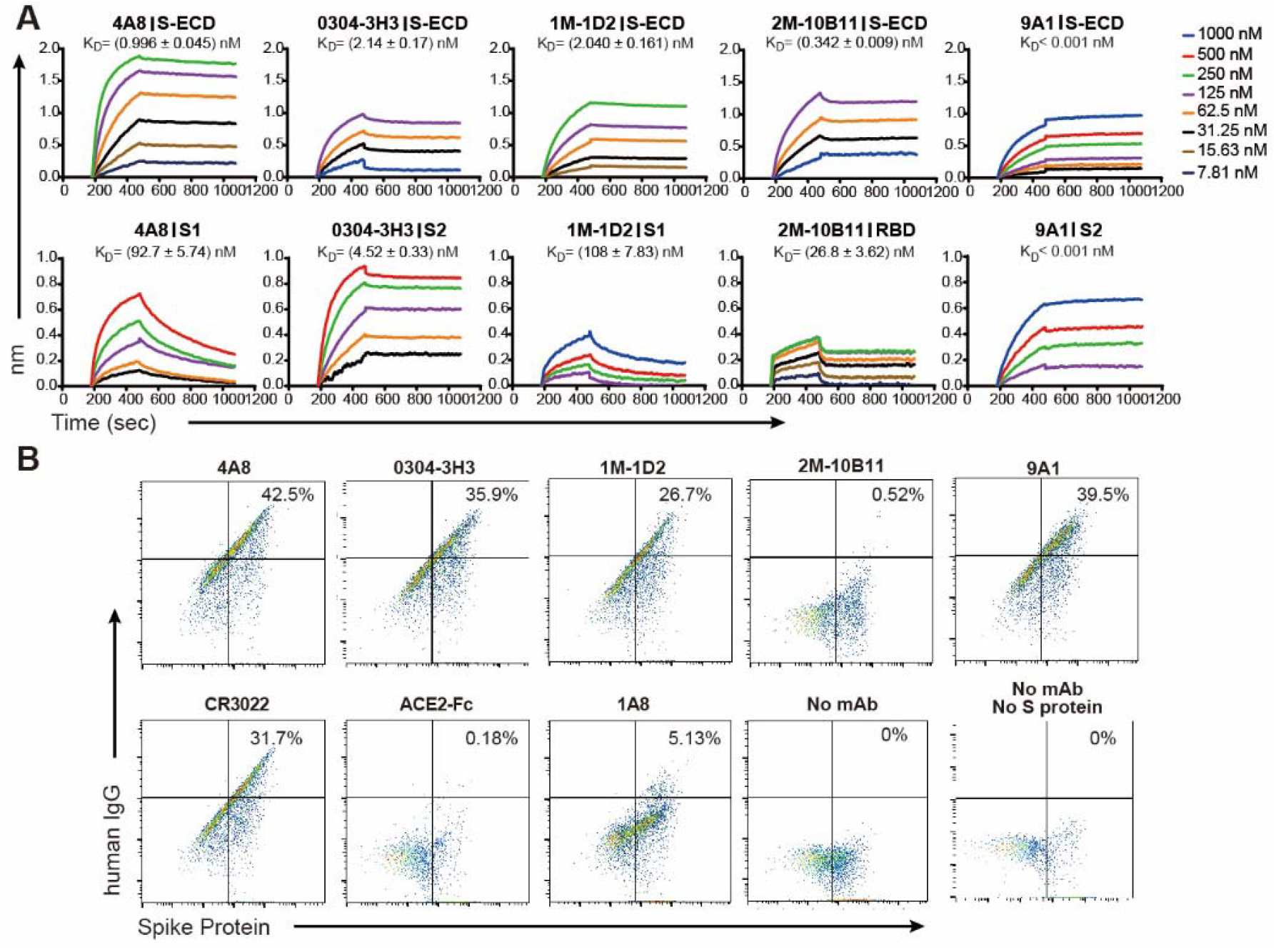
4A8 did not block the binding of Spike protein to ACE2 receptor. **(A)** BLI sensorgrams and kinetics of mAbs binding to S proteins. **(B)** The binding of S protein to human ACE2 overexpressing 293T cells were determined by flow cytometry. Following the preincubation of S protein with each indicated mAb, the mAb-S mixtures were added to the ACE2-expressing cells. cells were stained with anti-human IgG FITC (mAb binding, x-axis) and anti-His (S binding, y-axis). Percentages of double positive cells were shown. Control mAb CR3022 and 1A8 were previously reported to bind SARS-CoV RBD and Marburg glycoprotein, respectively, and ACE2-Fc protein was a human ACE2 protein conjugated with human Fc.

To further investigate whether these mAbs block the binding of S protein to ACE2, we performed flow cytometry using HEK 293T cells expressing human ACE2. As expected, only 10B11 among the 5 mAbs prevented S protein from binding to ACE2, with percentage of 0.52% for IgG and S protein double-positive cell (Fig 4B). However, CR3022, which competes with 10B11, did not interfere with the binding of S to ACE2 with double positive cell percentage of 31.7%. 4A8 also failed to interfere with binding of the S protein to ACE2.

### Cryo-EM structure of the complex between 4A8 and S-ECD

The mAb 4A8 was overexpressed and purified by Protein A resin from the B memory cells and S-ECD of SARS-CoV-2 was purified through M2 affinity resin and size exclusion chromatography. 4A8 and S-ECD protein were mixed and incubated at a stoichiometric ratio of ~ 1.2 to 1 for 1 hour and applied to size exclusion chromatography to remove excess proteins. The fraction containing the complex was concentrated for cryo-EM sample preparation.

To investigate the interactions between 4A8 and the S protein, we solved the cryo-EM structure of the complex at an overall resolution of 3.1 Å (Fig. 5, Movie S1). Details of cryo-EM sample preparation, data collection and processing, and model building can be found in Materials and Methods and supplementary materials (Figs. S3-S5). The S protein exhibits asymmetric conformations similar to the previously reported structures (*24, 25*), with one RBD “up” and the other two “down” (Fig. 5, fig. S3I).

**Fig. 5.**
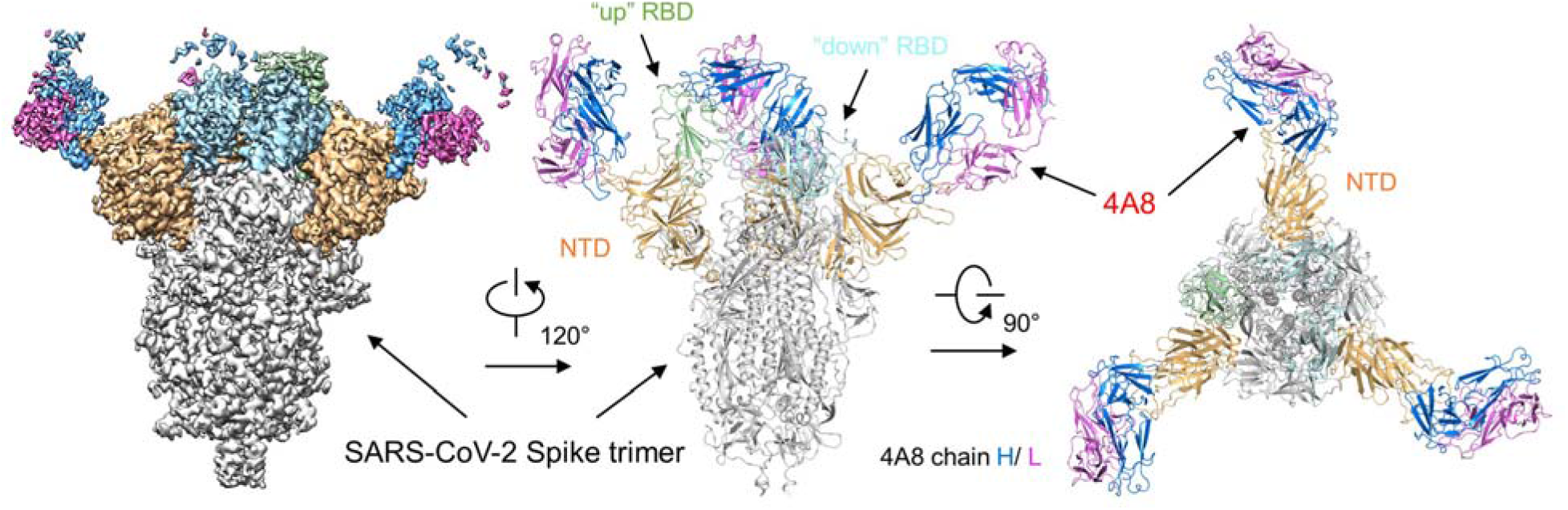
Cryo-EM structure of the 4A8 and S-ECD complex. The domain-colored cryo-EM map of the complex is shown on the right, and two perpendicular views of the overall structure are shown on the right. The heavy and light chains of 4A8 are colored blue and magenta, respectively. The NTDs of the trimeric S protein are colored orange.The one “up” RBD and two “down” RBDs of trimeric S protein are colored green and cyan, respectively.

### Recognition of the NTD by 4A8

Three 4A8 molecules bind to one trimeric S protein, each interacting with one NTD domain. Despite the different conformations of the three S protein protomers, the interface between 4A8 and each NTD is identical (Fig. 5). The map quality at the NTD-4A8 region was improved by focused refinement with a local resolution of 3.4 Å, enabling reliable analysis of the interactions between the NTD and the 4A8.

Association with 4A8 appears to stabilize the NTD epitope, which is invisible in the reported S protein structure alone (*24, 25*). Supported by the high resolution of NTD, we were able to build the structural model for three new loops for NTD, designated N1 (residues 67-79), N2 (residues 141-156), and N3 (residues 246-260), among which N2 and N3 loops mediate the interaction with 4A8.

Only the heavy chain of 4A8 participates in binding to the NTD mainly through three loops, named L1, L2 and L3 (Fig. 6A). The interface is constituted by extensive hydrophilic interaction network. R246 on the N3 loop of the NTD represents one docking site, simultaneously interacting with Glu1, Tyr27 and Glu31 of 4A8 (Fig. 6B). On the N2 loop of the NTD, Lys150 and Lys147 respectively form salt bridge with Glu54 and Glu72 of 4A8 (Fig. 6C). Lys150 is also hydrogen bonded (H-bond) with 4A8-Tyr111, while His146 forms a H-bond with 4A8-Thr30 (Fig. 6C). In addition to the hydrophilic interactions, Trp152 and Tyr145 on the N2 loop of the NTD also interact with Val102, Pro106, and Phe109 on the L3 loop of the mAb through hydrophobic and/or π-π interactions (Fig. 6D).

**Fig. 6.**
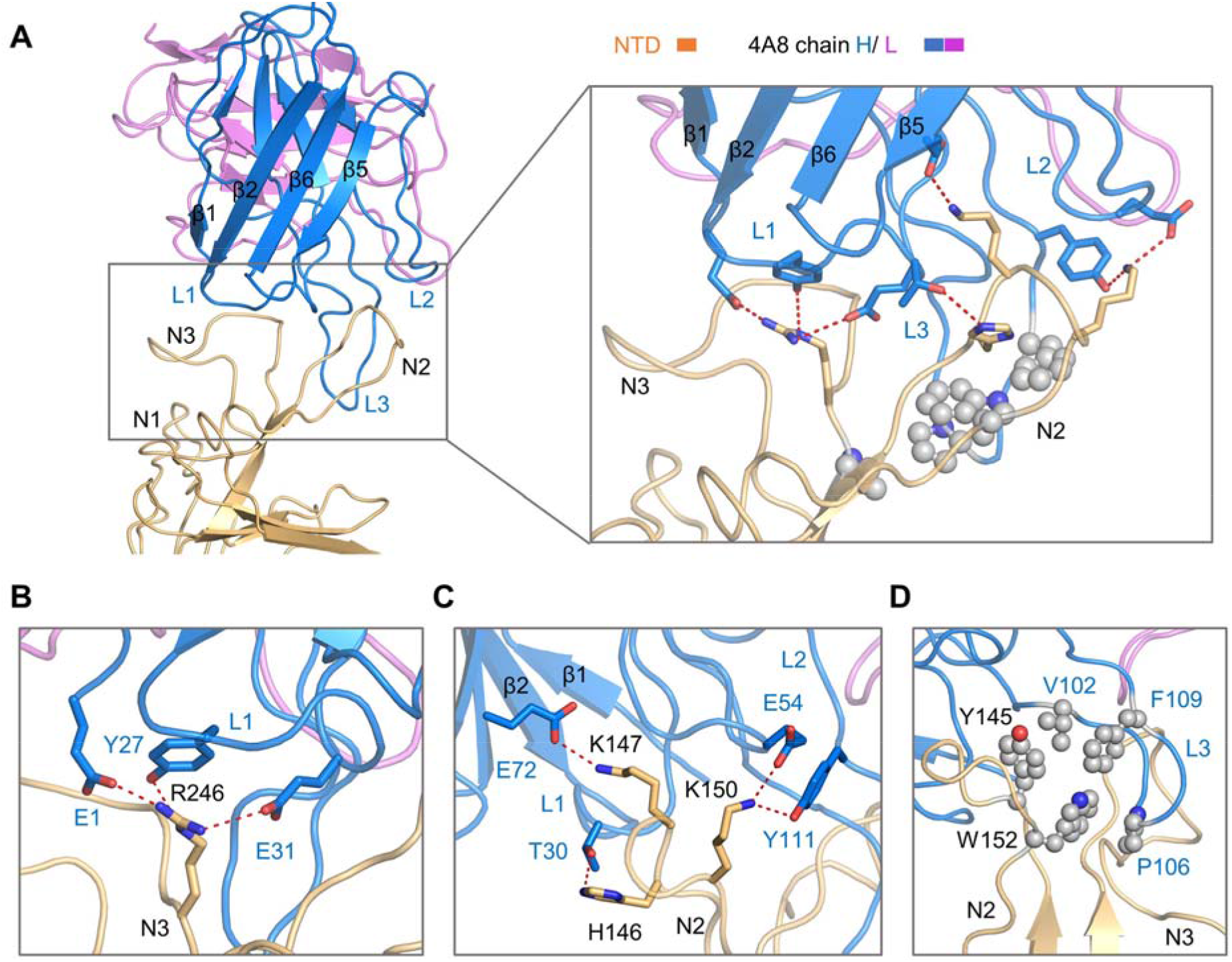
Interactions between the NTD and 4A8. **(A)** Extensive hydrophilic interactions on the interface between NTD and 4A8. Only one NTD-4A8 is shown. **(B-D)** Detailed analysis of the interface between NTD and 4A8. Polar interactions are indicated by red, dashed lines. The residues involved in hydrophobic interactions are presented as spheres.

## Discussion

There is an urgent need for prophylactic and therapeutic interventions for SARS-CoV-2 infections given the ongoing COVID-19 pandemic. No fully human neutralizing Abs have been reported, and little is known about the structural determinants of neutralization on which to base the rational selection of antibodies. Our work reveals that naturally occurring human SARS-CoV-2 mAbs isolated from the B cells of 10 recovered donors are diverse in gene usage and epitope recognition of S protein. Remarkably, the majority of the isolated mAbs did not recognize the RBD, and all the mAbs that neutralize live SARS-CoV-2 failed to inhibit the binding of S protein to ACE2. These unexpected results suggest the presence of other important mechanisms for SARS-CoV-2 neutralization in addition to suppressing the viral interaction with the receptor.

The S1-targeting mAb 4A8 does not block the interaction between ACE2 and S protein, but exhibits high levels of neutralization against both authentic and pseudotyped SARS-CoV-2 *in vitro*. Many neutralizing antibodies against the SARS-CoV-2 were reported to target the RBD of the S protein and block the binding between RBD and ACE2. Our results show that 4A8 bind to the NTD of S protein with potent neutralizing activity. Previous study showed that mAb 7D10 could bind to the NTD of S protein of MERS-CoV probably by inhibiting the RBD-ACE2 binding and the prefusion to postfusion conformational change of S protein (*29*). We aligned the crystal structure of 7D10 in complex with the NTD of S protein of MERS-CoV with our complex structure and found that the interfaces between the mAb and the NTDs are partially overlapped (Fig. S6). 7D10 may inhibit the interaction between MERS-CoV and ACE2 through its light chain that is close to the RBD. In our complex, the light chain of 4A8 is away from the RBD (Fig. S6). Therefore, we speculate that 4A8 may neutralize SARS-CoV-2 by restraining the conformational changes of the S protein. Furthermore, sequences alignment of the S proteins from SARS-CoV-2, SARS-CoV, and MERS-CoV revealed varied NTD surface sequences that are respectively recognized by different mAbs (Fig. S7).

Overall, this work reports a fully human neutralizing mAb recognizing a vulnerable epitope of NTD on S protein of SARS-CoV-2, functioning with a mechanism that is independent of receptor binding inhibition. Combination of 4A8 with RBD-targeting antibodies may avoid the escaping mutations of virus and serve as promising “cocktail” therapeutics. The information obtained from these studies can be used for development of the structure-based vaccine design against SARS-CoV-2.

## Acknowledgments

We thank the Cryo-EM Facility and Supercomputer Center of Westlake University for providing cryo-EM and computation support, respectively.

## Funding

This work was funded by the National Key R&D Program of China (2020YFC0841400), the National Natural Science Foundation of China (projects 31971123, 81803429, 81703048, 31900671, 81920108015, 31930059), the Key R&D Program of Zhejiang Province (2020C04001), the SARS-CoV-2 emergency project of the Science and Technology Department of Zhejiang Province (2020C03129) and the Leading Innovative and Entrepreneur Team Introduction Program of Hangzhou.

## Author contributions

W.C., Q.Z. and J.L. conceived the project. X.C., R.Y., J.Z., G.Z., Y.Z., Y.G., Y.L., L.X., M.H., Z.Z., P.F., Y.D., Z.C., J.L.Z., X.S., Y.C., L.F., L.H., J.X. and C.Y. did the experiments. All authors contributed to data analysis. X.C., R.Y., J.L., Q.Z. and W.C. wrote the manuscript.

## Competing interests

Authors declare no competing interests.

## Data and materials availability

Atomic coordinates and cryo EM density maps of the S protein of SARS-CoV-2 in complex bound with 4A8 (PDB: 7C2L; whole map: EMD-30276, antibody-epitope interface-focused refined map: EMD-30277) have been deposited to the Protein Data Bank (http://www.rcsb.org) and the Electron Microscopy Data Bank (https://www.ebi.ac.uk/pdbe/emdb/), respectively.

## Supplementary Materials

Materials and Methods

Figures S1-S7

Tables S1-S3

Movie S1

## Notes

### Competing Interest Statement

The authors have declared no competing interest.

## References and Notes

1. N. Zhu et al., A Novel Coronavirus from Patients with Pneumonia in China, 2019. N Engl J Med, (2020).

2. P. Zhou et al., A pneumonia outbreak associated with a new coronavirus of probable bat origin. Nature, (2020).

3. N. Zhu et al., A Novel Coronavirus from Patients with Pneumonia in China, 2019. N Engl J Med 382, 727–733 (2020).

4. F. Di Pierro, A. Bertuccioli, I. Cavecchia, Possible therapeutic role of a highly standardized mixture of active compounds derived from cultured Lentinula edodes mycelia (AHCC) in patients infected with 2019 novel coronavirus. Minerva gastroenterologica e dietologica, (2020).

5. Q. Gao et al., Rapid development of an inactivated vaccine candidate for SARS-CoV-2. Science, (2020).

6. C. Huang et al., Clinical features of patients infected with 2019 novel coronavirus in Wuhan, China. Lancet 395, 497–506 (2020).

7. T. G. Ksiazek et al., A novel coronavirus associated with severe acute respiratory syndrome. N Engl J Med 348, 1953–1966 (2003).

8. A. M. Zaki, S. van Boheemen, T. M. Bestebroer, A. D. Osterhaus, R. A. Fouchier, Isolation of a novel coronavirus from a man with pneumonia in Saudi Arabia. N Engl J Med 367, 1814–1820 (2012).

9. W. H. Li et al., Angiotensin-converting enzyme 2 is a functional receptor for the SARS coronavirus. Nature 426, 450–454 (2003).

10. J. H. Kuhn, W. Li, H. Choe, M. Farzan, Angiotensin-converting enzyme 2: a functional receptor for SARS coronavirus. Cell. Mol. Life Sci. 61, 2738–2743 (2004).

11. K. Kuba et al., A crucial role of angiotensin converting enzyme 2 (ACE2) in SARS coronavirus-induced lung injury. Nat. Med. 11, 875–879 (2005).

12. D. S. Dimitrov, The secret life of ACE2 as a receptor for the SARS virus. Cell 115, 652–653 (2003).

13. J. Lan et al., Structure of the SARS-CoV-2 spike receptor-binding domain bound to the ACE2 receptor. Nature, (2020).

14. Q. Wang et al., Structural and Functional Basis of SARS-CoV-2 Entry by Using Human ACE2. Cell, (2020).

15. R. Yan et al., Structural basis for the recognition of the SARS-CoV-2 by full-length human ACE2. Science, (2020).

16. J. Shang et al., Structural basis of receptor recognition by SARS-CoV-2. Nature, (2020).

17. M. Hoffmann et al., SARS-CoV-2 Cell Entry Depends on ACE2 and TMPRSS2 and Is Blocked by a Clinically Proven Protease Inhibitor. Cell, (2020).

18. M. Hoffmann et al., SARS-CoV-2 Cell Entry Depends on ACE2 and TMPRSS2 and Is Blocked by a Clinically Proven Protease Inhibitor. Cell 181, 271–280 e278 (2020).

19. P.-H. Wang, Y. Cheng, Increasing Host Cellular Receptor—Angiotensin-Converting Enzyme 2 (ACE2) Expression by Coronavirus may Facilitate 2019-nCoV Infection. bioRxiv, (2020).

20. T. M. Gallagher, M. J. Buchmeier, Coronavirus spike proteins in viral entry and pathogenesis. Virology 279, 371–374 (2001).

21. G. Simmons, P. Zmora, S. Gierer, A. Heurich, S. Pohlmann, Proteolytic activation of the SARS-coronavirus spike protein: cutting enzymes at the cutting edge of antiviral research. Antiviral Res 100, 605–614 (2013).

22. S. Belouzard, V. C. Chu, G. R. Whittaker, Activation of the SARS coronavirus spike protein via sequential proteolytic cleavage at two distinct sites. Proc Natl Acad Sci U S A 106, 5871–5876 (2009).

23. W. Song, M. Gui, X. Wang, Y. Xiang, Cryo-EM structure of the SARS coronavirus spike glycoprotein in complex with its host cell receptor ACE2. PLoS Pathog 14, e1007236 (2018).

24. D. Wrapp et al., Cryo-EM structure of the 2019-nCoV spike in the prefusion conformation. Science, (2020).

25. A. C. Walls et al., Structure, Function, and Antigenicity of the SARS-CoV-2 Spike Glycoprotein. Cell 181, 281–292 e286 (2020).

26. C. Krempl, B. Schultze, H. Laude, G. Herrler, Point mutations in the S protein connect the sialic acid binding activity with the enteropathogenicity of transmissible gastroenteritis coronavirus. J Virol 71, 3285–3287 (1997).

27. F. Kunkel, G. Herrler, Structural and functional analysis of the surface protein of human coronavirus OC43. Virology 195, 195–202 (1993).

28. G. Lu, Q. Wang, G. F. Gao, Bat-to-human: spike features determining ‘host jump’ of coronaviruses SARS-CoV, MERS-CoV, and beyond. Trends Microbiol 23, 468–478 (2015).

29. H. Zhou et al., Structural definition of a neutralization epitope on the N-terminal domain of MERS-CoV spike glycoprotein. Nat Commun 10, 3068 (2019).

30. B. Ju et al., Potent human neutralizing antibodies elicited by SARS-CoV-2 infection. bioRxiv, (2020).

31. C. Wang et al., A human monoclonal antibody blocking SARS-CoV-2 infection. Nat Commun 11, 2251 (2020).

32. D. Wrapp et al., Structural Basis for Potent Neutralization of Betacoronaviruses by Single-Domain Camelid Antibodies. Cell, (2020).

33. X. Chen et al., Human monoclonal antibodies block the binding of SARS-CoV-2 spike protein to angiotensin converting enzyme 2 receptor. Cell Mol Immunol, (2020).

34. M. Yuan et al., A highly conserved cryptic epitope in the receptor-binding domains of SARS-CoV-2 and SARS-CoV. Science, (2020).

35. C. G. Price, P. W. Abrahams, Copper tolerance in a population of Silene vulgaris ssp. maritima (A. & D. Love) at Dolfrwynog Bog near Dolgellau, North Wales. Environ Geochem Health 16, 27–30 (1994).

