## Supplementary Materials for "A potent neutralizing human antibody reveals the N-terminal domain of the Spike protein of SARS-CoV-2 as a site of vulnerability"

**This PDF file includes:**

Materials and Methods

Figs. S1 to S7

Table S1 to S3

Movie. S1

Supplemental References

### Materials and methods

#### Ethic Statement

A written informed consent was regularly obtained from all patients. The study was approved by the Ethics Committee of Wuhan Infectious Disease Hospital, Hubei Province, China.

#### Viruses and cells culture

SARS-CoV-2 was isolated from the lung lavage fluid of an infected patient. All work with infectious SARS-CoV-2 was performed in the biosafety level 3 (BSL-3) facility of Beijing Institute of Microbiology and Epidemiology, Academy of Military Medical Sciences, China. Vero E6 cells, HEK293T cells and ACE2-293T cells were grown in Dulbecco's modified Eagle's medium (DMEM) (Gibco) supplemented with 10% fetal bovine serum (Gibco), penicillin (100 IU/mL) and streptomycin (100 IU/mL). ACE2-293T cells, which are HEK293T cells expressing ACE2 receptor, were produced and kept in our laboratory. Expi293F cells (Thermo Scientific) were grown in Expi293™ Expression Medium (Thermo Scientific). HEK293F cells were grown in SMM 293T-II medium (Sino Biological Inc.).

#### Human subjects

After informed consent was obtained, peripheral blood samples were collected from ten survivors of SARS-CoV-2. Peripheral blood mononuclear cells (PBMCs) were isolated using human lymphocyte separation medium (Dakewe Biotech) according to the manufacturer's protocol. Briefly, peripheral blood was diluted with phosphate buffered saline (PBS) (Gibco). The mixture was then slowly transferred above lymphocyte separation medium. The volume of blood, PBS and the separation medium were the same. After centrifuging at room temperature at 800 g for 20 minutes, PBMCs were separated and collected to a new centrifuge tube. Following two steps of washing with PBS, the PBMCs were resuspended with Cell Freezing Medium (Thermo Scientific) and stored at -80C° until use.

#### Fluorescent Cell sorting and single-cell PCR

To sort antibody secreting cells (ASCs), PBMCs were stained with anti-CD3-PerCP (BD), anti-CD20-PerCP (BD), anti-CD19-Alexa Fluor700 (Beckman Coulter), anti-CD27-PE-Cy7 (Beckman Coulter), and anti-CD38-FITC (Stem Cell). ASCs were gated as $CD19^{+}CD3^{-}CD20^{-}CD27^{High}CD38^{High}$. To sort SARS-CoV-2 S protein specific memory B cells, PBMCs were stained with anti-CD3-PerCP (BD), anti-CD19-Alexa Fluor700 (Beckman Coulter), anti-IgG-PE (BD), and SARS-CoV-2 S protein (His tag) (Sino Biological), followed by washing with PBS and staining with anti-His tag-Alexa Fluor647 (Thermo Scientific). The SARS-CoV-2 S protein specific memory B cells were defined as $CD3^{-}CD19^{+}IgG^{+}S\text{-}ECD^{+}$. Single gated cells were sorted into 96-well plates (Bio-Rad) containing 20 µL RNase-free-water (TransGen Biotech) and 20 U RNase inhibitor (Promega) in each well. Double-stranded cDNA was produced using One-Step reverse transcription PCR (RT-PCR) kit (Qiagen). Antibody VH and VL genes were amplified through nested PCR using TransStart Taq DNA polymerase (TransGen Biotech) as described previously (*1*). The PCR products were sequenced by Sangon Biotech. Sequences were analyzed using IMGT/ V-QUEST (http://www.imgt.org/IMGT_vquest).

**Protein expression and purification**

The full-length Ig heavy- and light-chain linear expression cassettes were produced by assembling the cytomegalovirus (CMV) promoter, Ig leader sequence, VH/VL gene, Ig constant region (IgG1), and poly(A) sequence fragment together using overlapping PCR as previously described (*2*). The antibody VH/VL and constant region genes were then amplified and cloned into expression vector pcDNA3.4 using NEBuilder® HiFi DNA Assembly Cloning Kit (NEB). The plasmids of paired Ig H and L genes were co-transfected into Expi293F expression system (Thermo Scientific) following the manufacturer's protocol to produce recombinant mAbs. HiTrap rProtein A column (GE Healthcare) was used to purify antibodies from cell culture supernatants.

The extracellular domain (ECD) (1-1208 a.a) was cloned into the pCAG vector (Invitrogen) with two proline substitutions at residues 986 and 987, a "GSAS" substitution at residues 682 to 685 and a C-terminal T4 fibritin trimerization motif followed by one Flag tag. The mutants were generated with a standard two-step PCR-based strategy.

The recombinant S-ECD protein was overexpressed using the HEK 293F mammalian cells (Invitrogen) at 37°C under 5% $CO_2$ in a Multitron-Pro shaker (Infors, 130 rpm). When the cell density reached 2.0 $\times 10^6$ cells/mL, the plasmid was transiently transfected into the cells. To transfect one liter of cell culture, about 1.5 mg of the plasmid was premixed with 3 mg of polyethylenimines (PEIs) (Polysciences) in 50 mL of fresh medium for 15 mins before adding to cell culture. Cells were removed by centrifugation at 4000×g for 15 mins after sixty hours transfection. The secreted S-ECD proteins were purified by anti-FLAG M2 affinity resin (Sigma Aldrich). After loading two times, the anti-FLAG M2 resin was washed with the wash buffer containing 25 mM Tris (pH 8.0), 150 mM NaCl. The protein was eluted with the wash buffer plus 0.2 mg/mL flag peptide. The eluent was then concentrated and subjected to size-exclusion chromatography (Superose 6 Increase 10/300 GL, GE Healthcare) in buffer containing 25 mM Tris (pH 8.0), 150 mM NaCl. The peak fractions were collected and concentrated to incubate with mAb 4A8. The purified S-ECD was mixed with the mAb 4A8 at a molar ratio of about 1:1.2 for one hour. Then the mixture was subjected to size-exclusion chromatography (Superose 6 Increase 10/300 GL, GE Healthcare) in buffer containing 25 mM Tris (pH 8.0), 150 mM NaCl. The peak fractions were collected for EM analysis.

**Enzyme-linked immunosorbent assay (ELISA)**

Polystyrene microplates (Corning) were coated overnight with 2 μg/mL SARS-CoV-2 S, S1, S2 or RBD protein (Sino Biological). After washing with PBS containing 0.2% Tween 20 (Solarbio Life Sciences), the plates were blocked using 2% BSA (Sigma Aldrich) in PBST for 1 h at 37C°. Following washing with PBST, serial dilutions of testing antibodies were added to each well and incubated at 37C° for 1 h. After washing with PBST, horseradish peroxidase (HRP)-conjugated anti-human IgG antibody (Abcam) was added at the dilution of 1:10000 and incubated at 37C° for 1 h. After washing, TMB single-component substrate solution (Solarbio Life Sciences) was added to the microplate and incubated at room temperature for 6 min, followed by adding 2M $H_2SO_4$ to stop the reaction. The absorbance was detected at 450 nm/630 nm. The data was analyzed using GraphPad Prism 7.0.

**Competition binding analysis**

Detecting antibodies were conjugated with biotin using EZ-Link™ Sulfo-NHS-Biotin (Thermo Scientific) following the manufacturer' s protocol. Microplates were coated with 2 μg/mL SARS-CoV-2 S protein (Sino Biological) overnight at 4 ℃°. Detecting antibodies at the final concentration of 1 μg/mL and blocking antibodies at the final concentration of 100 μg/mL were added to the same well and incubated for 1 h at 37 °C. After washing with PBST, streptavidin conjugated with HRP (Thermo Scientific) was added at the concentration 1 μg/mL and incubated for 1 h at 37 °C. Following washing with PBST, TMB single-component substrate solution (Solarbio Life Sciences) was added and incubated at room temperature for 6 min. 2 M $H_2SO_4$ was then added to stop the reaction. The absorbance was detected at 450nm/630nm. The competition value was determined by comparing the binding value of detecting antibody in the presence of blocking antibody divided by the binding value of detecting antibody alone.

**Sequence analysis of antibody variable region genes**

Antibody sequences were annotated with igBLAST (*3*) and IMGT/V-QUEST (*4*) and then trimmed to extract only the variable region from FWR1 to the end of the J gene. Based on the gene annotation results of igBLAST, the count and relative abundance of V(D)J alleles, genes or families were determined by R package 'Alakazam' in Change-O toolkit (*5*). R package 'ggpubr' and 'ggplot2' were used for pie chart. Circos plots indicating the combination of V/D/J alleles were generated by self-written Perl scripts and R scripts. R package 'circlize' was used for data visualization. Antibody amino acid sequence alignments and evolutionary analyses were conducted in MEGA7 (*6*). The evolutionary history was inferred using the Neighbor-Joining method (*7*). The bootstrap consensus tree inferred from 500 replicates (*8*) was taken to represent the evolutionary history of the taxa analyzed (*8*). Branches corresponding to partitions reproduced in less than 50% bootstrap replicates were collapsed. The evolutionary distances were computed using the Maximum Composite Likelihood method (*9*). To analyze sequence similarity, antibody DNA sequences were compared to each other (pairwise alignments) by ClustalW (*10*), a multiple alignment program. The scores were calculated from separate pairwise alignments by the method of Wilbur and Lipman method (*11*) with the default DNA weight matrix (IUB). Data format conversion were completed by self-written Perl scripts and R scripts. R package 'corrplot' was used for data visualization.

**Neutralization experiment**

Vero-E6 cells were inoculated in 96-well cell culture plates (20,000 cells per well) with DMEM (Gibco) supplemented with 10% fetal bovine serum and grown overnight at 37 ℃. Sera or antibodies were serially diluted and then mixed with 100 $TCID_{50}$ SARS-CoV-2. The mixture was moved to the wells containing Vero-E6 cells and incubated at 37 ℃ for 1 h. Following removing the supernatants, 200 μL cell culture medium were added and the plates were then incubated at 37C° with 5% $CO_2$ for 3 days. Cells were stained with crystal violet and absorbance at 570nm/630nm were measured. Neutralization was defined as percentage reduction compared to positive controls. Neutralization titers of two replicates were calculated using a non-linear regression analysis in GraphPad Prism 7.

Supernatants at 3 days post infection were harvested and tested using quantitative real-time PCR to determine the number of viral genomes. RNA extraction was performed using RNAeasy Plus mini kit (Qiagen) following the manufacturer's protocol. Eluted RNA was used for reverse transcription using Maxima First Strand cDNA Synthesis Kit for RT-qPCR (Thermo Scientific) according to the manufacturer's instructions with oligo(dT). Subsequently, quantitative real-time PCR was performed using TaqMan Universal Master Mix Ⅱ with UNG (Thermo Scientific) according to the manufacturer's instructions, with the primer CoV-RT-F3 5'- GGGGAACTTCTCCTGCTAGAAT-3', primer CoV-RT-R3 5'- CAGACATTTTGCTCTCAAGCTG-3' and a probe CoV-RT-P3 5'- FAM-TTGCTGCTGCTTGACAGATT-3'. The primers and probe were purchased from Sangon Biotech. The $IC_{50}$ was calculated using a non-linear regression analysis in GraphPad Prism 7.

**Pseudotyped virus packaging and neutralizing**

Gene encoding full-length SARS-CoV-2 S protein (Genebank ID: QHD43416.1) was inserted into the pDC316 vector, resulting in plasmid pDC316- SARS-CoV-2-S. A total of $7.0 \times 10^6$ ACE2-293T cells were inoculated in a 10 cm cell culture dish and grown overnight at 37C° with 5% $CO_2$. The pDC316-SARS-CoV-2-S and HIV backbone vector pNL4-3.Luc.R-E- were co-transfected into 293T cells with Lipofectamine3000 transfection reagent (Invitrogen). The culture medium was replaced after 6 hours. The supernatants containing HIV-pseudotyped virus with S protein were collected 48 h post-transfection and filtered through a 0.45 μm filter. The supernatants were then aliquoted and stored at -80 °C. To determine the neutralization ability of sera or antibodies, pseudotyped virus was incubated with serially diluted sera or antibodies at 37 °C for 1 h and then added to 96-well culture plates containing $2 \times 10^4$ ACE2-293T cells with two replicates. The cells were then maintained at 37 °C with 5% $CO_2$ for 48 h. The cells were lysed and luciferase activity was measured using firefly luciferase assay system (Promega). Neutralization was calculated as (1-Lucmeasured/Lucvirus control) × 100%. Neutralization titers were calculated using a non-linear regression analysis in GraphPad Prism 7.

**Biolayer interferometry**

Antibodies to be tested were diluted to the concentration of 5 μg/mL with PBS containing 0.05% Tween 20 (Solarbio Life Sciences) and then immobilized onto Anti-hIgG Fc Capture (AHC) biosensors (Sartorius AG). After a 60-seconds washing step with PBST, biosensor tips were immersed into the wells containing SARS-CoV-2 S-ECD/S1/S2/RBD protein (Sino Biological) of serial dilutions and allowed to associate for 300 seconds, followed by a dissociation step of 600 seconds. The Kd value for each pair of antibody and antigen was calculated using Data Analysis Software 9.0 (Sartorius AG).

**Flow cytometry receptor binding inhibition assay**

Antibodies to be tested were incubated at the concentration of 400 μg/mL with 40 μg/mL SARS-CoV-2 S protein (Sino Biological) in 50 μL PBS for 1 h at 37 C°. The mAb-S protein mixture was then incubated with $5 \times 10^5$ ACE2-293T cells for 30 min at room temperature. After washing twice with PBS containing 2% FBS (Gibco), cells were resuspended and incubated with anti-His tag antibody conjugated with Alexa Fluor 647 (Thermo Scientific) and anti-human IgG antibody conjugated with FITC (Abcam) for 30 min at room temperature. Cells were washed twice and resuspended with PBS containing 2% FBS before analyzed using BD FACSCanto II. ACE2-Fc (Novoprotein) was used as positive control. Data in FCS format were analyzed using FlowJo X 10.0.7r2.

**Cryo-EM sample preparation**

The peak fractions of S-ECD and mAb 4A8 complex was concentrated to about 1.5 mg/mL and applied to the grids. Aliquots (3.3 μL) of the protein complex were placed on glow-discharged holey carbon grids (Quantifoil Au R1.2/1.3). The grids were blotted for 2.5 s or 3.0 s and flash-frozen in liquid ethane cooled by liquid nitrogen with Vitrobot (Mark IV, Thermo Scientific). The cryo-EM samples were transferred to a Titan Krios operating at 300 kV equipped with Cs corrector, Gatan K3 Summit detector and GIF Quantum energy filter. Movie stacks were automatically collected using AutoEMation (*12*), with a slit width of 20 eV on the energy filter and a defocus range from -1.2 µm to -2.2 µm in super-resolution mode at a nominal magnification of 81,000×. Each stack was exposed for 2.56 s with an exposure time of 0.08 s per frame, resulting in a total of 32 frames per stack. The total dose rate was approximately 50 $e^{-}/Å^{2}$ for each stack. The stacks were motion corrected with MotionCor2 (*13*) and binned 2-fold, resulting in a pixel size of 1.087 Å/pixel. Meanwhile, dose weighting was performed (*14*). The defocus values were estimated with Gctf (*15*).

**Data processing**

Particles were automatically picked using Relion 3.0.6 (*16-19*) from manually selected micrographs. After 2D classification with Relion, good particles were selected and subject to one cycle of heterogeneous refinement without symmetry using cryoSPARC (*20*).The good particles were selected and subjected to homogeneous refinement with C1 symmetry, resulting in the 3D reconstruction for the whole structures, which was further subject to 3D auto-refinement and post-processing with Relion. For interface between SARS-CoV-2 S protein and mAb 4A8, the dataset were subject to focused refinement with adapted mask on each NTD-4A8 sub-complex to improve the map quality. Then the dataset of three NTD-4A8 sub-complexes were combined and subject to focused refinement with Relion, resulting in the 3D reconstruction of better quality on the interface between S protein and mAb 4A8.

The resolution was estimated with the gold-standard Fourier shell correlation 0.143 criterion (*21*) with high-resolution noise substitution (*22*). Refer to Supplemental Figures S4-S5 and Supplemental Table S3 for details of data collection and processing.

**Model building and structure refinement**

For model building of the complex of S protein of SARS-CoV-2 with mAb 4A8, the atomic model of S protein (PDB ID: 6VSB) and an mAb molecule (PDB ID: 5FUZ) were used as templates, which were molecular dynamics flexible fitted (*23*) into the whole cryo-EM map of the complex and the focused-refined cryo-EM map of the NTD-4A8 sub-complex, respectively. And the fitted atomic models were further manually adjusted with Coot (*24*). Each residue was manually checked with the chemical properties taken into consideration during model building. Several segments, whose corresponding densities were invisible, were not modeled. Structural refinement was performed in Phenix (*25*) with secondary structure and geometry restraints to prevent overfitting. To monitor the potential overfitting, the model was refined against one of the two independent half maps from the gold-standard 3D refinement approach. Then, the refined model was tested against the other map. Statistics associated with data collection, 3D reconstruction and model building were summarized in Table S3.

### Supplementary Figures and Legends

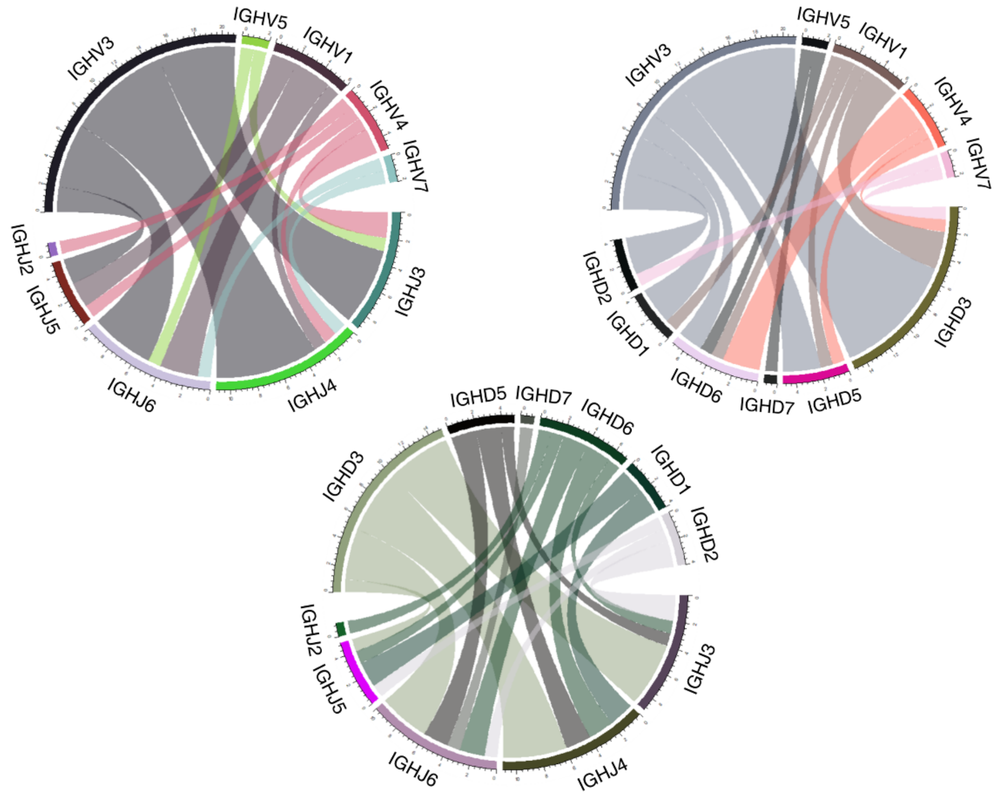


**Fig. S1**

Combination of heavy chain V/D/J genes of the thirty-five spike protein-specific antibodies. Each arc of the circle represents a family of IGH V/D/J gene.

The bands between two families represent the combination of V and D, V and J or D and J genes. The width of bands represents the frequency of combination.

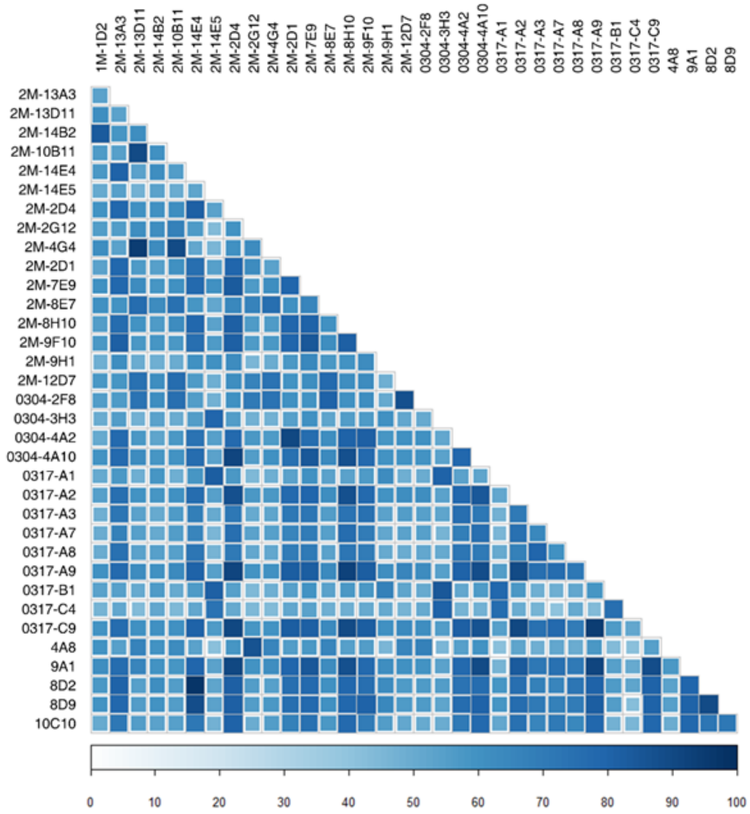


**Fig. S2**

Quantification of CDRH3 sequence identities of the thirty-five spike protein-specific antibodies by similarity index.

The index was determined using BLOSUM (Blocks Substitution Matrix).

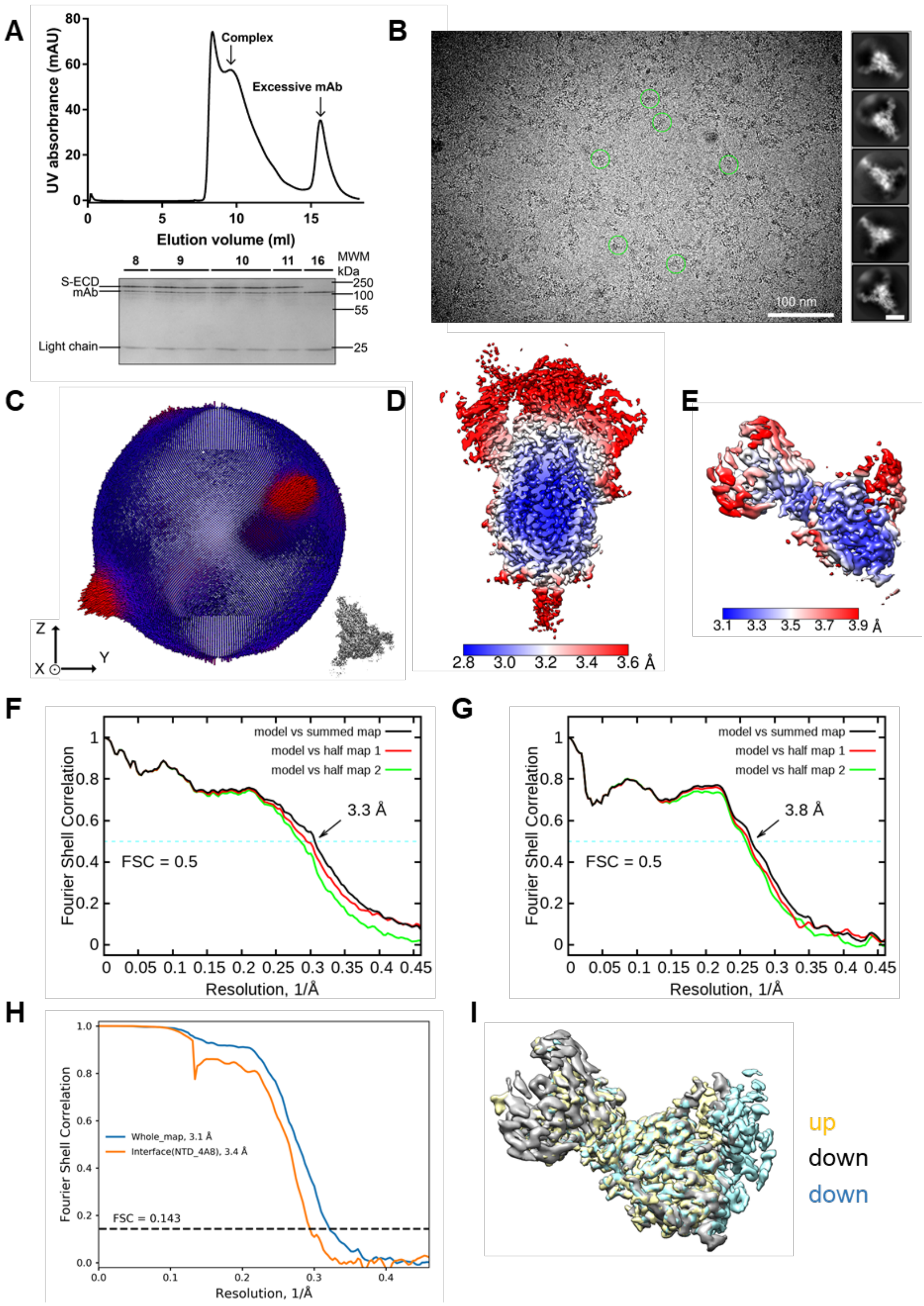


**Fig. S3**

Cryo-EM analysis of S-ECD of SARS-CoV-2 bound with 4A8 complex.

**(A)** Representative SEC purification profile of the S-ECD of SARS-CoV-2 in complex with mAb 4A8. **(B)** Representative cryo-EM micrograph and 2D class averages of cryo-

EM particle images. The scale bar in 2D class averages is 10 nm. **(C)** Euler angle distribution in the final 3D reconstruction of S-ECD of SARS-CoV-2 bound with 4A8 complex. **(D)** and **(E)** Local resolution maps for the 3D reconstruction of the overall structure and interface between NTD of S-ECD and 4A8, respectively. **(F)** FSC curve of the refined model of S-ECD of SARS-CoV-2 bound with 4A8 complex versus the overall structure that it is refined against (black); of the model refined against the first half map versus the same map (red); and of the model refined against the first half map versus the second half map (green). The small difference between the red and green curves indicates that the refinement of the atomic coordinates is not enough overfitting. **(G)** FSC curve of the refined model of interface between NTD of S-ECD and 4A8, which is the same as the (F). **(H)** FSC curve of the overall structure (blue) and interface between NTD of S-ECD and 4A8 (orange). **(I)** Superposition in local map of interface between NTD of S-ECD and 4A8 for different conformation of SARS-CoV-2 S protein trimer in "up" or "down" RBD, which has no difference among three maps. The maps for one "up" RBD and two "down" RBDs are colored yellow and cyan, respectively.

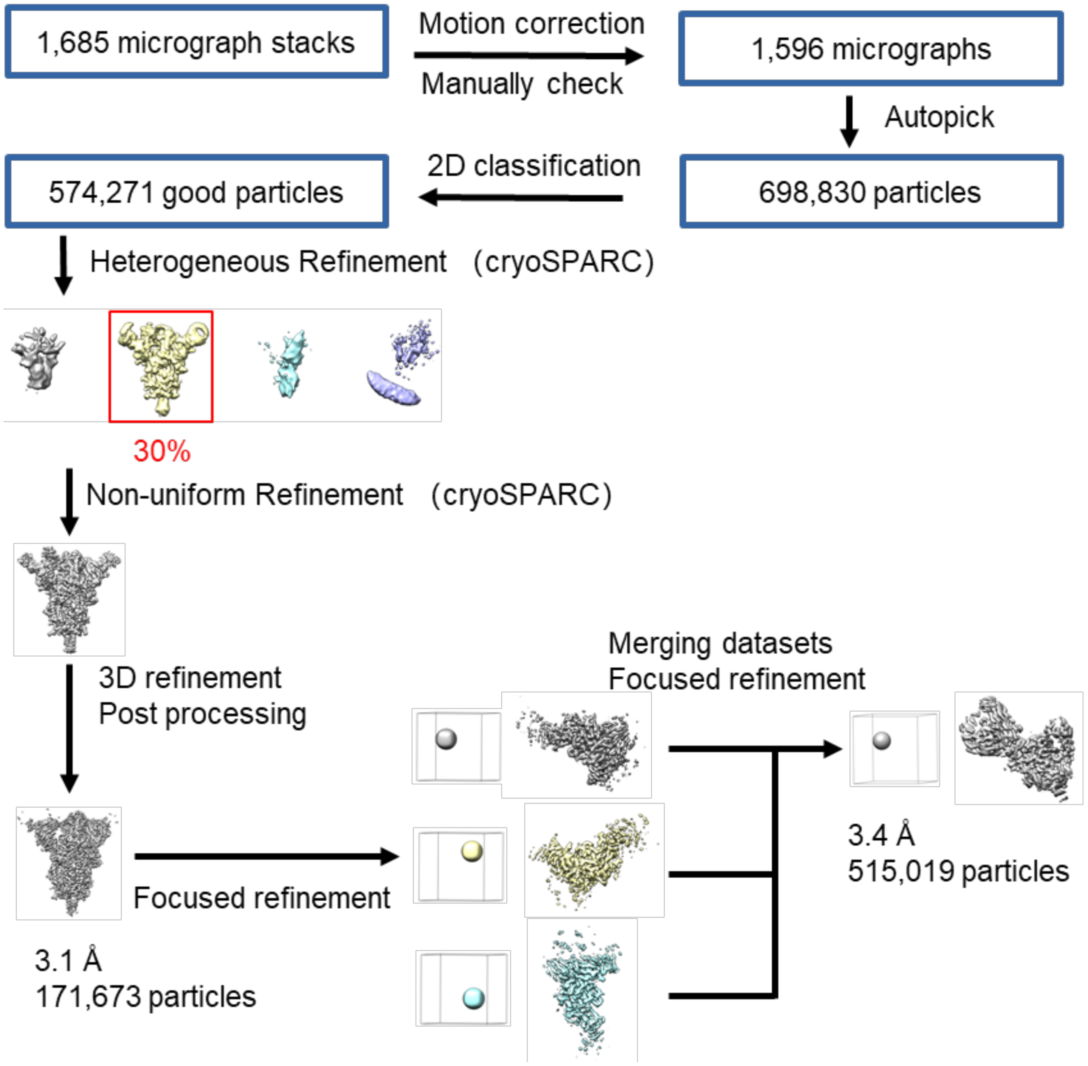


**Fig. S4**

Flowchart for cryo-EM data processing of S-ECD of SARS-CoV-2 bound with 4A8 complex.

Please refer to the ‘Data Processing’ section in Methods for details.

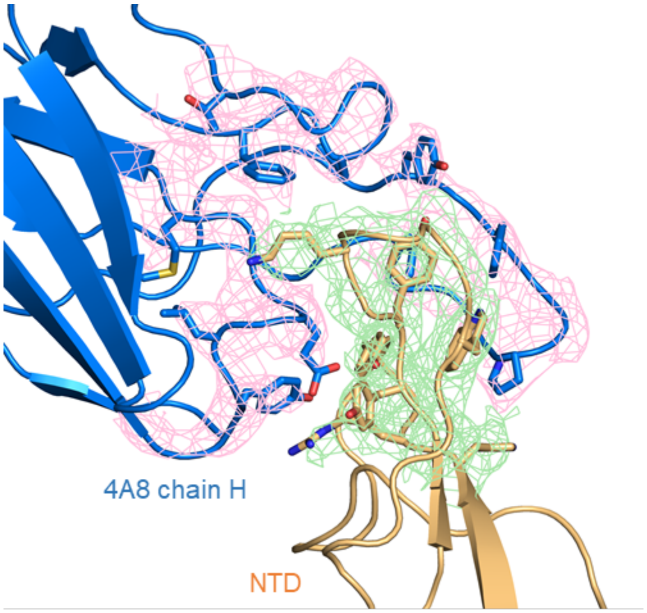


**Fig. S5**

Cryo-EM density of the interface between NTD of S-ECD and 4A8.

The density is contoured at 12 σ, which shown as pink meshes in chain H of 4A8 and green meshes in NTD of S-ECD, respectively.

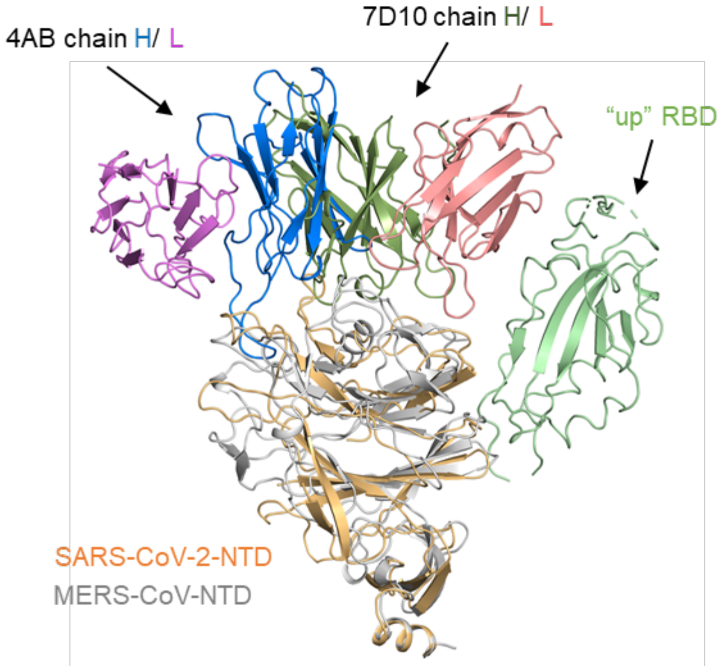


**Fig. S6**

Structural comparison of SASR-CoV-2 bound with 4A8 complex and MERS-CoV bound with 7D10 complex (PDB ID: 6J11).

Shown here is the NTD of MERS-CoV superimposing with the NTD of SASR-CoV-2, the NTD can be aligned with root mean squared deviation of ~ 8.6 Å over ~ 193 pairs of Cα atoms, which is overlapping in heavy chain between 4A8 and 7D10.

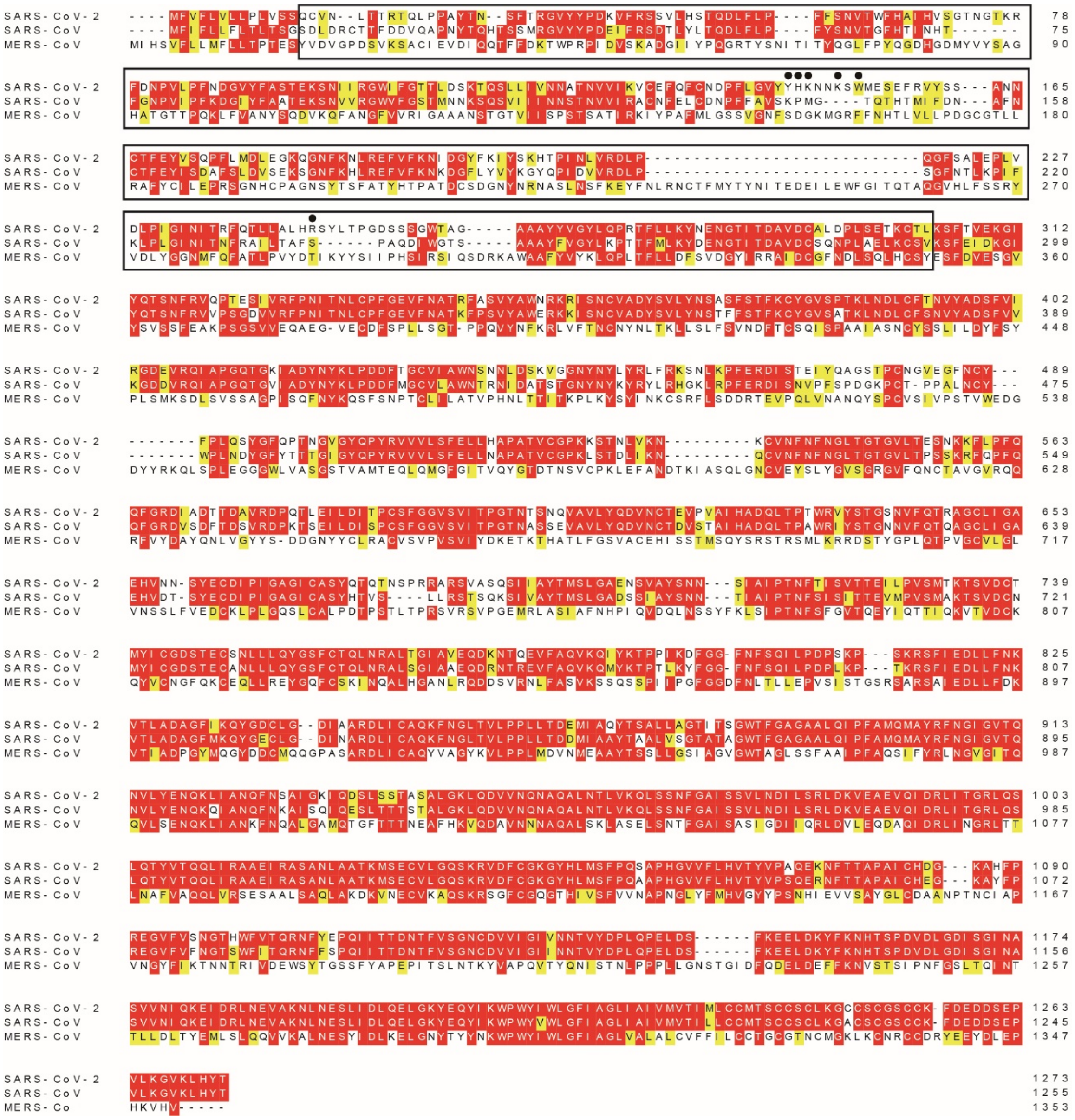

**Fig. S7**

Sequence alignment for S protein of SARS-CoV-2， SARS-CoV and MERS-CoV.

The sequences were aligned using ClustalX. The aligned sequences are S protein of SARS-CoV-2, SARS-CoV and MERS-CoV. Amino acids that are identical or conserved in at least 2 sequences are colored red or yellow. The amino acid residues located in the N-terminal domain are lined out by black box. Key residues involved in interaction with mAb 4A8 are marked with black circles. Uniprot IDs are listed below: **S protein of SARS-CoV-2**: P0DTC2; **S protein of SARS-CoV**: P59594; **S protein of MERS-CoV**: K9N5Q8.

**Table S1**

**Clinical characteristics of COVID-19 recovered patients**

| ID | Age (Years) | Gender | Duration from blood collection date to disease confirmation date (Days) | First Symptoms |
|---|---|---|---|---|
| Pt 1 | 43 | M | 29 | Cough, fever, and sore muscle |
| Pt 2 | 25 | M | 29 | Fever and fatigue |
| Pt 3 | 30 | M | 31 | Fever and chest tightness |
| Pt 4 | 35 | F | 29 | Fever |
| Pt 5 | 50 | M | 23 | Fever and cough |
| Pt 6 | 32 | F | 11 | Fever, cough, and fatigue |
| Pt 7 | 53 | M | 15 | Fever and cough |
| Pt 8 | 28 | F | 15 | Fever and sore muscle |
| Pt 9 | 43 | M | 12 | Fever |
| Pt 10 | 36 | F | 10 | Fever |

**Table S2**

**Origination, germline sequence and variable region CDR3 sequence of the 35 S protein -specific antibodies.**

| Sequence ID | Donor | IGHV | VH-CDR3 | IGLV | VL-CDR3 |
|---|---|---|---|---|---|
| 1M-1D2 | 1 | IGHV3-64*01 | ARGAEYYDFWSG YYSAYFDY | IGLV1-47*01 | SMGAQPDLC A |
| 2M-13A3 | 2 | IGHV3-30*04 | ARGGGSYYYWFD P | IGKV4-1*01 | QQYYST |
| 2M-13D11 | 2 | IGHV4-34*01 | ARAGYSSSWYGV RGVDP | IGKV3-20*01 | QQYGSSRSWT |
| 2M-14B2 | 2 | IGHV3-30*18 | AKGSDIVVVPVGN WFDP | IGKV1-39*01 | QQSYSTFTLY T |
| 2M-10B11 | 2 | IGHV3-66*02 | ARATWLRGVMDV | IGLV6-57*02 | QSYDSSNHW V |
| 2M-14E4 | 2 | IGHV4-61*01 | ARVQRYYPDSSGF YGRRFDI | IGKV1-39*01 | QQSHSFPFT |
| 2M-14E5 | 2 | IGHV3-30*04 | ARSGGGSYRGPFD Y | IGKV1-9*01 | QQLNSYVT |
| 2M-2D4 | 2 | IGHV3-23*04 | AKIGLGLGGLLRR YFDY | IGKV3-11*01 | QQRTNWPL |
| 2M-2G12 | 2 | IGHV3-11*04 | DANYDYVVAQTG YGR | IGKV1-39*01 | QQNYSTWT |
| 2M-4G4 | 2 | IGHV1-46*01 | ARERGDSSGYYEII TTANRRFGMDV | IGLV2-23*01 | CSYAVSSTWV |
| 2M-2D1 | 2 | IGHV4-39*01 | ARGDRIQLWLLDA FDI | IGLV1-47*01 | CSTGAQPEWL G |
| 2M-7E9 | 2 | IGHV1-69*01 | ARIPGWDRGTDRN WNDD | IGKV3-11*01 | QQRSNWPPAF T |
| 2M-8E7 | 2 | IGHV1-69*01 | ARTYSFDSSGYYY DY | IGKV3-11*01 | QQHSNWPPKI T |
| 2M-8H10 | 2 | IGHV3-30*04 | ARAFYDSNWSVGS YFDS | IGKV4-1*01 | QQYYNNQWT |
| 2M-9F10 | 2 | IGHV3-9*01 | AKDSVRREYTHAR VPFDN | IGKV1-39*01 | QQSFVSPRT |
| 2M-9H1 | 2 | IGHV3-30*18 | AKSSKIFYLGESRE VDY | IGKV1-17*01 | LQHKSYPLT |
| 2M-12D7 | 2 | IGHV3-9*01 | AKDVRYCSSTSCY FSAFDI | IGKV1-39*01 | QQSYSTPRT |
| 0304-2F8 | 3 | IGHV5-51*01 | ARRGDGLYYYGM DV | IGKV2-28*01 | MQALQTPQT |
| 0304-3H3 | 4 | IGHV4-59*01 | ARDRIAPVGKFFG WYFDL | IGKV3-15*01 | QQYNKWPPW T |

| | | | | | |
|---|---|---|---|---|---|
| 0304-4A2 | 4 | IGHV4-39*07 | ARELFTAVAGKGG IDY | IGKV4-1*01 | HQYYNTPRT |
| 0304-4A10 | 4 | IGHV3-64*01 | ARSSSRGFDY | IGKV4-1*01 | QQYYSSPYA |
| 0317-A1 | 5 | IGHV3-30*18 | AKDFKGGSSSWYT PEIEYGMDV | IGKV3-11*01 | QQRSNWPPT |
| 0317-A2 | 5 | IGHV7-4-1*02 | ARLIRHEAHTYCS GGSCYSPDYYYG MDV | IGKV1-39*01 | QQSYSTPPT |
| 0317-A3 | 5 | IGHV3-48*03 | ASNPPLGEPYFDI | IGKV1-39*01 | QQTYRPPWT |
| 0317-A7 | 5 | IGHV3-30-3*01 | ARWGGGMQYLDV | IGKV2-28*01 | MQTLQTPYT |
| 0317-A8 | 5 | IGHV7-4-1*02 | ARAGPNYDFWSG YYQTFDY | IGKV1-33*01 | QQYDNLPLT |
| 0317-A9 | 5 | IGHV1-24*01 | ATATAMDGYYYY YGMDV | IGKV2-24*01 | MQATQFPYT |
| 0317-B1 | 5 | IGHV5-51*01 | ASAGSSWYGDAFD I | IGKV4-1*01 | QQYYSTYGS |
| 0317-C4 | 5 | IGHV1-24*01 | ATATIFGVANNWF DP | IGLV1-51*01 | GTWDSSLSVV V |
| 0317-C9 | 5 | IGHV3-30*18 | AKDLGYYDILTGQ LGGYYYYYGMDV | IGLV1-44*01 | AAWDDSLNG VV |
| 4A8 | 6/7/8/9/ 10 | IGHV1-24*01 | ATSTAVAGTPDLF DYYYGMDV | IGKV2-24*01 | TQATQFPYT |
| 9A1 | 6/7/8/9/ 10 | IGHV3-30*18 | AKVSAIFWLGQGL SPIDV | IGKV3-15*01 | HQYSKWPVT |
| 8D2 | 6/7/8/9/ 10 | IGHV3-7*01 | ARDWDYDILTGS WFGAFDI | IGKV1-17*01 | LQHNSYPLT |
| 8D9 | 6/7/8/9/ 10 | IGHV3-7*01 | ARPTIGYSYGSDY | IGLV3-1*01 | QAWDSSTGV |
| 10C10 | 6/7/8/9/ 10 | IGHV3-7*01 | ARDWDYDILTGS WFGAFDI | IGKV1-17*01 | LQHNNYPLT |

**Table S3**

**Cryo-EM data collection and refinement statistics.**

| | | |
|---|---|---|
| **Data collection** | | |
| EM equipment | Titan Krios (Thermo Fisher Scientific) | |
| Voltage (kV) | 300 | |
| Detector | Gatan K3 Summit | |
| Energy filter | Gatan GIF Quantum, 20 eV slit | |
| Pixel size (Å) | 1.087 | |
| Electron dose (e-/Å2) | 50 | |
| Defocus range (μm) | -1.2 ~ -2.2 | |
| Number of collected micrographs | 1,685 | |
| Number of selected micrographs | 1,596 | |
| Sample | S-ECD of SARS-CoV-2 bound with 4A8 complex | |
| **3D Reconstruction** | | |
| | Overall | Interface between NTD and 4A8 |
| Software | cryoSPARC/ Relion | Relion |
| Number of used particles | 171,673 | 515,019 |
| Resolution (Å) | 3.1 | 3.4 |
| Symmetry | C1 | |
| Map sharpening B factor ($Å^2$) | -90 | |
| **Refinement** | | |
| Software | Phenix | |
| Cell dimensions (Å) | 313.056 | |
| Model composition | | |
| Protein residues | 4,420 | |
| Side chains assigned | 4,420 | |
| Sugar | 74 | |
| R.m.s deviations | | |
| Bonds length (Å) | 0.008 | |
| Bonds Angle (˚) | 1.032 | |
| Ramachandran plot statistics (%) | | |
| Preferred | 90.98 | |
| Allowed | 8.62 | |
| Outlier | 0.40 | |

**Movie S1.**
**Cryo-EM structure of the 4A8 and S-ECD complex revealing the recognition of the NTD by monoclonal antibody.**

**Supplementary references:**


1. K. Smith *et al.*, Rapid generation of fully human monoclonal antibodies specific to a vaccinating antigen. *Nat Protoc* **4**, 372-384 (2009).
2. H. X. Liao *et al.*, High-throughput isolation of immunoglobulin genes from single human B cells and expression as monoclonal antibodies. *J Virol Methods* **158**, 171-179 (2009).
3. J. Ye, N. Ma, T. L. Madden, J. M. Ostell, IgBLAST: an immunoglobulin variable domain sequence analysis tool. *Nucleic Acids Res* **41**, W34-40 (2013).
4. V. Giudicelli, X. Brochet, M. P. Lefranc, IMGT/V-QUEST: IMGT standardized analysis of the immunoglobulin (IG) and T cell receptor (TR) nucleotide sequences. *Cold Spring Harb Protoc* **2011**, 695-715 (2011).
5. N. T. Gupta *et al.*, Change-O: a toolkit for analyzing large-scale B cell immunoglobulin repertoire sequencing data. *Bioinformatics* **31**, 3356-3358 (2015).
6. S. Kumar, G. Stecher, K. Tamura, MEGA7: Molecular Evolutionary Genetics Analysis Version 7.0 for Bigger Datasets. *Mol Biol Evol* **33**, 1870-1874 (2016).
7. N. Saitou, M. Nei, The neighbor-joining method: a new method for reconstructing phylogenetic trees. *Mol Biol Evol* **4**, 406-425 (1987).
8. J. Felsenstein, CONFIDENCE LIMITS ON PHYLOGENIES: AN APPROACH USING THE BOOTSTRAP. *Evolution* **39**, 783-791 (1985).
9. K. Tamura, M. Nei, S. Kumar, Prospects for inferring very large phylogenies by using the neighbor-joining method. *Proc Natl Acad Sci U S A* **101**, 11030-11035 (2004).
10. J. D. Thompson, T. J. Gibson, D. G. Higgins, Multiple sequence alignment using ClustalW and ClustalX. *Curr Protoc Bioinformatics* **Chapter 2**, Unit 2.3 (2002).
11. W. J. Wilbur, D. J. Lipman, Rapid similarity searches of nucleic acid and protein data banks. *Proc Natl Acad Sci U S A* **80**, 726-730 (1983).
12. J. Lei, J. Frank, Automated acquisition of cryo-electron micrographs for single particle reconstruction on an FEI Tecnai electron microscope. *Journal of structural biology* **150**, 69-80 (2005).
13. S. Q. Zheng *et al.*, MotionCor2: anisotropic correction of beam-induced motion for improved cryo-electron microscopy. *Nature methods* **14**, 331-332 (2017).
14. T. Grant, N. Grigorieff, Measuring the optimal exposure for single particle cryo-EM using a 2.6 A reconstruction of rotavirus VP6. *eLife* **4**, e06980 (2015).
15. K. Zhang, Gctf: Real-time CTF determination and correction. *Journal of structural biology* **193**, 1-12 (2016).
16. J. Zivanov *et al.*, New tools for automated high-resolution cryo-EM structure determination in RELION-3. *eLife* **7**, (2018).
17. D. Kimanius, B. O. Forsberg, S. H. Scheres, E. Lindahl, Accelerated cryo-EM structure determination with parallelisation using GPUs in RELION-2. *eLife* **5**, (2016).
18. S. H. Scheres, RELION: implementation of a Bayesian approach to cryo-EM structure determination. *Journal of structural biology* **180**, 519-530 (2012).
19. S. H. Scheres, A Bayesian view on cryo-EM structure determination. *Journal of molecular biology* **415**, 406-418 (2012).

20. A. Punjani, J. L. Rubinstein, D. J. Fleet, M. A. Brubaker, cryoSPARC: algorithms for rapid unsupervised cryo-EM structure determination. *Nature methods* **14**, 290-296 (2017).
21. P. B. Rosenthal, R. Henderson, Optimal determination of particle orientation, absolute hand, and contrast loss in single-particle electron cryomicroscopy. *Journal of molecular biology* **333**, 721-745 (2003).
22. S. Chen *et al.*, High-resolution noise substitution to measure overfitting and validate resolution in 3D structure determination by single particle electron cryomicroscopy. *Ultramicroscopy* **135**, 24-35 (2013).
23. L. G. Trabuco, E. Villa, K. Mitra, J. Frank, K. Schulten, Flexible fitting of atomic structures into electron microscopy maps using molecular dynamics. *Structure (London, England : 1993)* **16**, 673-683 (2008).
24. P. Emsley, B. Lohkamp, W. G. Scott, K. Cowtan, Features and development of Coot. *Acta crystallographica. Section D, Biological crystallography* **66**, 486-501 (2010).
25. P. D. Adams *et al.*, PHENIX: a comprehensive Python-based system for macromolecular structure solution. *Acta crystallographica. Section D, Biological crystallography* **66**, 213-221 (2010).
